# Quantitative comparison of single-cell RNA sequencing versus single-molecule RNA imaging for quantifying transcriptional noise

**DOI:** 10.1101/2024.08.09.607289

**Authors:** Neha Khetan, Binyamin Zuckerman, Giuliana P. Calia, Xinyue Chen, Ximena Garcia Arceo, Leor S. Weinberger

**Affiliations:** Gladstone|UCSF Center for Cell Circuitry, University of California, San Francisco, CA 94158; Department of Biochemistry and Biophysics, University of California, San Francisco, CA 94158; Department of Pharmaceutical Chemistry, University of California, San Francisco, CA 94158

**Keywords:** transcriptional noise, noise-enhancer molecule, single-cell RNA sequencing (scRNA-seq), single molecule RNA FISH (smFISH)

## Abstract

Stochastic fluctuations (noise) in transcription generate substantial cell-to-cell variability. However, how best to quantify genome-wide noise, remains unclear. Here we utilize a small-molecule perturbation (IdU) to amplify noise and assess noise quantification from numerous scRNA-seq algorithms on human and mouse datasets, and then compare to noise quantification from single-molecule RNA FISH (smFISH) for a panel of representative genes. We find that various scRNA-seq analyses report amplified noise, without altered mean-expression levels, for ∼90% of genes and that smFISH analysis verifies noise amplification for the vast majority of genes tested. Collectively, the analyses suggest that most scRNA-seq algorithms are appropriate for quantifying noise including a simple normalization approach, although all of these systematically underestimate noise compared to smFISH. From a practical standpoint, this analysis argues that IdU is a globally penetrant noise-enhancer molecule—amplifying noise without altering mean-expression levels—which could enable investigations of the physiological impacts of transcriptional noise.

**MOTIVATION:** Cell-to-cell variability in isogenic populations is predominantly attributed to the stochastic fluctuations (i.e., noise) in transcription. However, the quantification of this noise, particularly on a genome-wide scale, remains an open question. To address this general question, here we utilize a small-molecule perturbation reported to amplify transcriptional noise. Previous single-cell RNA-sequencing (scRNA-seq) analysis indicated that the pyrimidine nucleobase 5′-iodo-2′-deoxyuridine (IdU) amplifies noise but technical drawbacks of scRNA-seq may have obscured the penetrance and degree of noise amplification. Consequently, here we assess numerous scRNA-seq algorithms, on two different scRNA-seq datasets for a human and mouse cell type, for their ability to quantify noise amplification and then compare the results to noise quantification from single-molecule RNA FISH (smFISH) imaging for a panel of representative genes. The specific questions addressed are whether IdU amplifies noise in a globally or partially penetrant manner, and what scRNA-seq algorithm is most appropriate for quantifying transcriptional noise.

Graphical Abstract

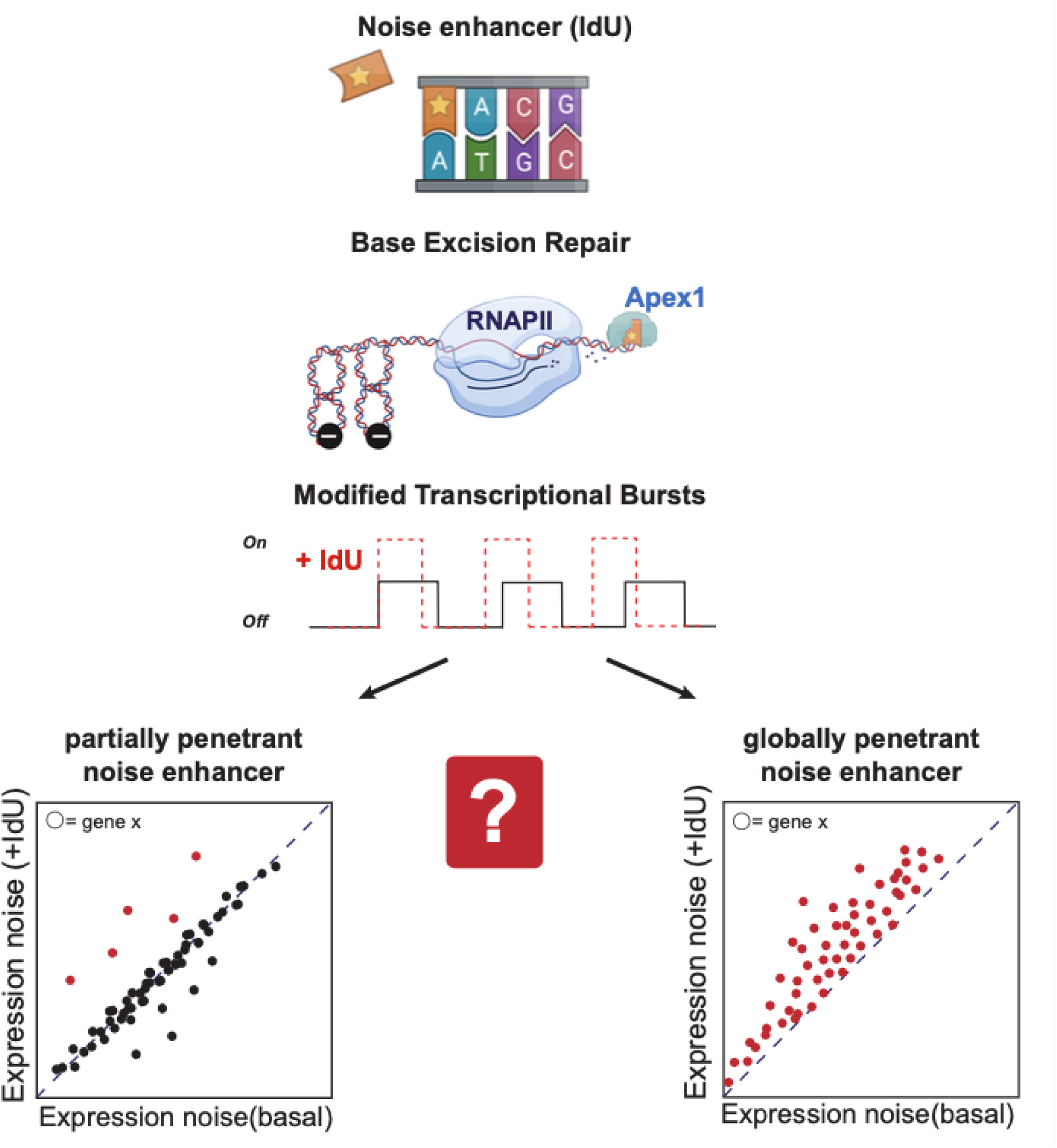

## INTRODUCTION

Cell-to-cell variability is an unavoidable consequence of the biochemical processes occurring in individual cells^1^ and has been implicated in cell-fate specification decisions ranging from HIV latency to cancer^2–4^. While a portion of cell-to-cell variability arises from extrinsic factors (e.g., cell size, cycle phase, or microenvironment), a substantial body of literature has demonstrated that, in isogenic populations of cells—particularly mammalian cells—a large fraction of the variability originates from intrinsic sources, such as stochastic fluctuations (noise) in transcription^5,6^. These intrinsic stochastic fluctuations can be quantitatively accounted by gene expression ‘toggling’ between active and inactive states which produces episodic ‘bursts’ of transcription, and a theoretical formalism commonly known as the two-state or random-telegraph model of gene expression is often used to fit these expression bursts^7–10^. Notably, transcriptional bursts are known to transmit to the protein level and expression noise is often amplified by nuclear export and mRNA processing in the cytoplasm^11,12^. Ultimately, these transcriptional bursts generate a substantial fraction of the measured noise^13,14^, and influence cell-fate specification decisions Nevertheless, measurements of transcriptional noise, particularly on a genome-wide scale, face several technical challenges. Specifically, how best to quantify noise across the transcriptome remains an open question, in part because there are no established methods to perturb noise across the transcriptome, which would be critical to benchmark noise-measurement approaches. Unfortunately, perturbing expression noise without altering the mean-expression level (i.e., orthogonal perturbation) has been challenging since for most physical processes the noise and mean levels are inherently linked. Intuitively, the reason for this linkage is that cellular processes, at their most fundamental level, are molecular birth-death processes and such birth-death processes are Poissonian (or often super-Poissonian) where the variance (σ^2^) necessarily equals the mean (μ) (i.e., as μ changes, so does σ^2^)—notably, the two-state model is an extension of the simple birth-death processes and generates super-Poisson distributions^11,12,14^ where the mean and noise remain tightly correlated but where the variance is always greater than the mean (σ^2^ > μ). Ultimately, the practical outcome of this intrinsic linkage is that the common metric for quantifying expression noise, the coefficient of variation (CV, which is the standard deviation normalized by the mean: σ/μ), necessarily decreases as the mean increases (CV ∝ 1/μ)^15^. Consequently, in some cases, the normalized variance, or Fano factor (σ^2^/μ) can be used to compare noise for processes with different mean values as Fano factor does not scale with the mean^16,17^. Regardless, perturbations that orthogonally modulate noise without changing the mean remain a challenge.

Historically, a mechanism for breaking the 1/μ dependence of CV and orthogonally modulating noise is via specific autoregulatory architectures (e.g., feedback and feedforward)^12,18^. However, more recently small molecules called “noise enhancers” were found to generate increased expression noise without altering the mean-expression level^19^ by a process known as homeostatic noise amplification^20^; a notable contrast to transcriptional activators, which increase mean-expression levels. We reported the molecular mechanism for one particular class of noise-enhancer molecule, pyrimidine-base analogs such as 5′-iodo-2′-deoxyuridine (IdU), and used scRNA-seq to show that IdU increases noise across the transcriptome. Unfortunately, scRNA-seq, despite its utility in measuring genome-wide expression, suffer from well-established issues of technical noise^21,22^ due to small inputs of RNA, varying sequencing depth, amplification bias, dropouts, and differences in capture ability^21,22^ that could obscure the quantification of IdU penetrance and IdU-mediated degree of noise enhancement.

Several algorithms and analysis pipelines have been developed to address the challenges of scRNA-seq analysis^23–25^. These algorithms include approaches to minimize biases from data-transformation, varying sequencing depth, dropouts, and incorporate methods to accurately delineate between technical and biological noise. However, in general, there is no consensus on the appropriate pipeline to quantify transcriptional noise and the choice of algorithm often depends on the biological system being studied and the specific scientific question being addressed.

Here, we set out to determine which scRNA-seq pipelines were appropriate for quantifying transcriptional noise using the noise-enhancer molecule IdU^20^ as a noise perturbation. Specifically, we also set out to determine if IdU acted as a globally or partially penetrant noise-enhancer molecule. We examined multiple scRNA-seq algorithms using a mouse embryonic stem cell (mESC) dataset with and without IdU treatment where each algorithm can account for experimental and sampling biases and are intended to minimize extrinsic and technical noise^26^. Each algorithm identified a different proportion of the genes exhibiting amplified noise as well as differences in the magnitude of noise amplification. A follow-up scRNA-seq profiling of human Jurkat T lymphocytes confirmed that IdU-mediated noise amplification is not restricted to the specific biology of mESCs. To validate scRNA-seq measurements, we then employed single-molecule RNA FISH (smFISH)—the gold standard for mRNA quantification due to its high sensitivity for mRNA detection^27^—to probe a panel of genes from across the transcriptome that span a wide array of expression levels and represent a range cellular functions. Collectively, these analyses indicate that IdU amplifies noise of most genes (globally penetrant) and could, in principle, be a candidate to probe the physiological roles of expression noise for diverse genes of interest.

## RESULTS

### Alternate scRNA-seq algorithms generate differing profiles of expression noise indicating noise amplification (ΔFano > 1) for ∼90% of expressed genes

To examine how different scRNA-seq normalization methods influence the quantification of IdU-mediated noise amplification, we employed commonly-used algorithms to analyze scRNA-seq data from IdU-treated versus DMSO-treated (control) mESCs^20^. This high-quality dataset consists of a few hundred deeply-sequenced cells (>60% sequencing saturation) and allows reliable noise quantification even for moderately-expressed genes. Yet, this experimental design is prone to technical noise, stemming from varying sequencing depth and the absence of biological replicates, which could compromise the evaluation of the actual IdU-mediated transcriptional noise amplification. To obtain a more rigorous measurement of noise enhancement penetrance, we compared noise quantification from a simple normalization approach i.e., normalized by sequencing depth (described in methods, and referred here as raw method) to the five established scRNA-seq algorithms (**Fig 1**): *SCTransform*^28^, *scran*^29^, *Linnorm*^30^, *BASiCS*^31,32^, and *SCnorm*^33^. *SCTransform* is a commonly used normalization method that employs negative binomial model, including regularization and variance stabilization steps. While, *scran* estimates cell-specific size factors for normalization by deconvolving pooled expression data from groups of cells. *Linnorm* utilizes homogenously expressed genes to estimate factors for transformation followed by variance stabilization. *SCnorm* groups genes based on count-depth relationships and uses quantile regression to generate normalization factors. While, BASiCS employ a hierarchical Bayesian-framework to simultaneously estimate model parameters for normalization factors, technical noise, and both Poissonian and super-Poissonian noise explicitly.

**Figure 1:**
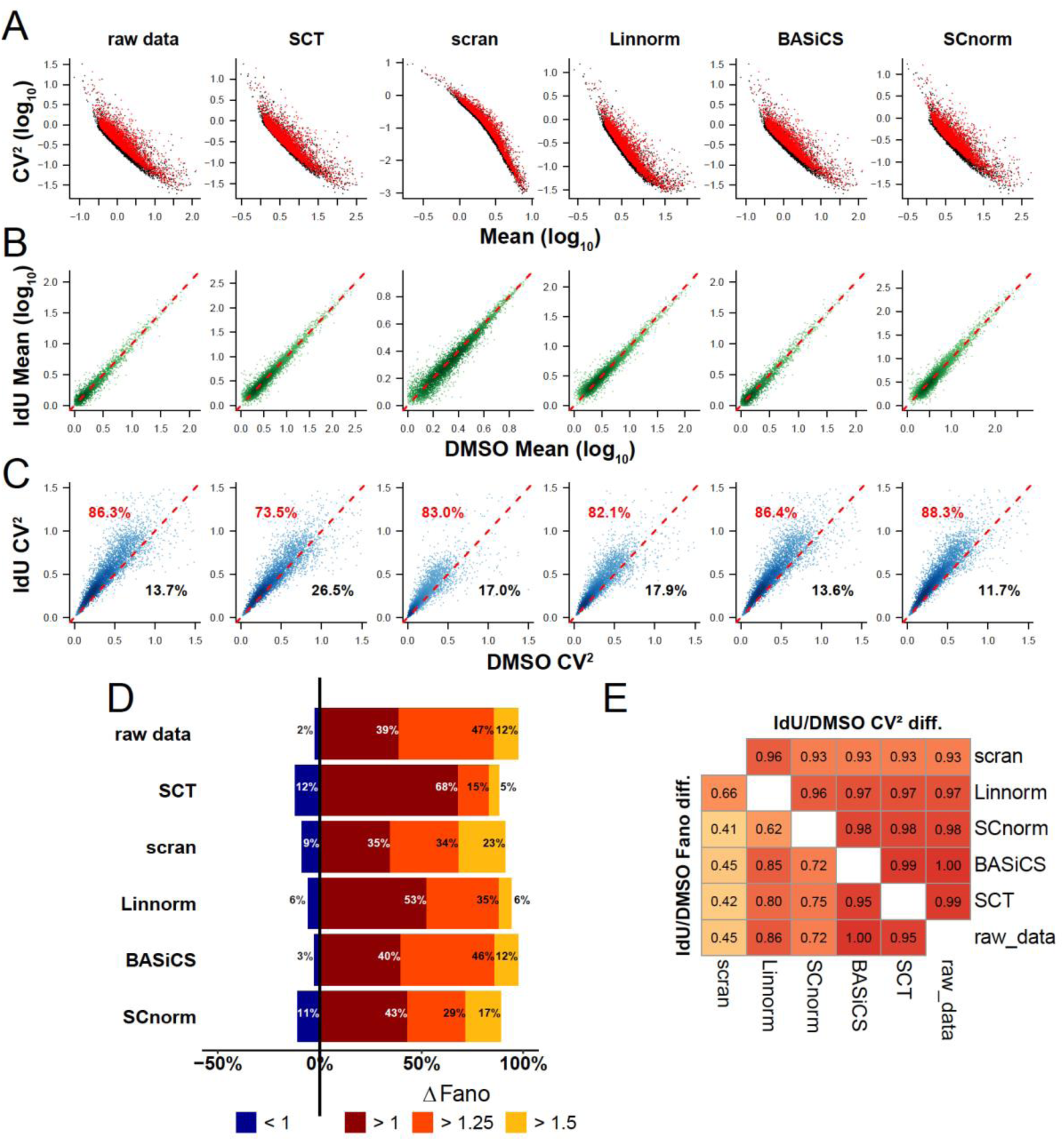
Common scRNA-seq normalization algorithms generate different quantifications of mRNA noise. **(A)** scRNA-seq analysis of CV^2^-vs-mean for 4,456 transcripts in mESCs treated with IdU (red) or DMSO control (black) as analyzed by commonly used normalization algorithms: SCTransform (“SCT”), scran, Linnorm, BASiCS, or SCnorm. **(B)** Mean expression for each of the 4,456 transcripts in presence and absence of IdU using each normalization algorithm; none of the normalizations algorithms generate substantial changes in mean expression for IdU-treated cells. **(C)** CV^2^ for each transcript in presence and absence of IdU using each normalization algorithm; different algorithms generate substantially different fractions of transcripts with amplified noise ranging from ∼70% of transcripts with amplified noise (SCTransform) to ∼88% of transcripts with amplified noise (SCnorm). **(D)** Quantification showing percentages of transcripts with indicated Fano factor fold changes between IdU-treated and control cells. **(E**) Pearson correlation coefficients between fold changes in noise metrics between IdU-treated and control cells. Shown are correlations between the indicated methods quantifying fold changes in CV^2^ (upper right heatmap) and Fano (lower left heatmap). See also Figure S1.

Despite their substantially different technical schemes, analysis with each of these normalizations indicated that IdU induces a substantial amplification of noise (CV^2^) for most expressed genes (**Fig. 1A**; Wilcoxon rank sum test for CV^2^: P<10^-17^ for all methods). The IdU-induced noise amplification appeared to be homeostatic (**Fig. 1B**), with mean-expression levels largely unchanged by IdU under all algorithms (Wilcoxon rank sum test for mean values: P > 0.02 for SCnorm and scran, and P > 0.1 for SCTransform, Linnorm and BASiCS). However, each algorithm calculated a somewhat different percentage of expressed genes with increased CV^2^ ranging from 73% to 88% of genes exhibiting increased noise (**Fig. 1C**). Quantification of noise by analysis of the normalized variance (i.e., Fano factor), which in principle eliminates the scaling dependence on mean-expression level, showed a similar profile **(Fig. 1D**; Wilcoxon rank sum test for Fano: P < 10^-70^ for all methods). Whilst every algorithm showed a majority of expressed genes exhibit amplified noise by Fano factor, each calculated a different penetrance (i.e., percentage of noise-amplified genes) as well as substantial differences in the magnitude of noise amplification among those transcripts with amplified noise.

Notably, the analysis further confirmed that BASiCS yielded minimal data-transformation when compared to the other algorithms (**Figs. S1 and S2**). As a negative control, the DMSO-treated samples were randomly split into two subpopulations and the noise metrics of both groups compared and no significant global change in CV^2^ or Fano factor between the two randomly split groups was found (**Fig. S2A-B**).

To explore the effect of IdU on transcriptional noise in a different biological context and under a different experimental setup, a separate scRNA-seq profiling +/- IdU was performed on second cell line: human Jurkat T lymphocytes. The Jurkat cell line (a suspension cell line) is more homogeneous than adherent stem cells^11^ and exhibits a cell-cycle time of about 24 h^34^, which is 2– 3x longer than mESCs^35^. IdU treatment duration and concentration were modified for Jurkat cells to account for the longer cell-doubling time and altered sensitivity and toxicity of Jurkats to IdU compared to mESCs, as determined by viability analysis under varying IdU doses (data not shown). Two biological replicates of Jurkat cells, each treated with DMSO or 20 uM IdU for 48h, were profiled by a moderate-depth scRNA-seq and comparison between the replicates confirmed that noise quantification by all algorithms is highly reproducible (**Fig. S2C)**, and that the batch-effect contribution to noise is negligible (**Fig. S2D**).

In this Jurkat scRNA-seq dataset, sequencing-depth bias was minimized and largely eliminated, since all samples and replicates exhibited highly uniform coverage as estimated by UMI counts per sample (**Fig. S2E**). Analysis with the different normalization methods indicated a consistent increase in noise metrics despite a lower fraction of affected genes compared to mESCs (**Fig. S3**). Overall, these results confirm that IdU enhances noise in diverse human and mouse cell types, and that the extent of noise generated depends (at least partly) on cell division rates (though other possible mechanisms may account for this difference, see Discussion).

### RNA quantification by smFISH indicates widespread penetrance of IdU-induced noise amplification

To directly quantify RNA levels in individual cells using a non-sequencing-based approach, we employed smFISH, a well-established imaging method to assess noise and the gold standard for quantitative assessment of mRNA expression^36^. While smFISH enables quantification of mRNA abundance in individual cells, it is a relatively low-throughput method requiring distinct fluorescent probes and image quantification for each transcript species of interest. Consequently, to generate a representative view of overall transcription using smFISH, we selected a panel of eight genes **(Fig. S4A-B)** that satisfied three criteria: (i) they displayed the greatest difference in noise amplification as calculated by the different RNA-seq algorithms (**Fig. S4B**); (ii) spanned a wide range of gene-expression levels (i.e., across 2-Logs) and genome locations (**Fig. S4A**)^37^; and (iii) the genes were compatible with smFISH probe design (i.e., a minimum of 30 probes were predicted to hybridize to the transcript; see **Data 3**). We also included previously reported smFISH data^20^ from a ninth gene, *Nanog*, to benchmark the smFISH analysis for the effect of IdU induced amplification of noise. smFISH probe sets for each gene in the panel were generated, cells were imaged in the presence/absence of IdU, and images (**Fig. 2A, Fig. S4C**) were segmented and analyzed using FishQuant^38^ to obtain per-cell mRNA counts.

**Figure 2:**
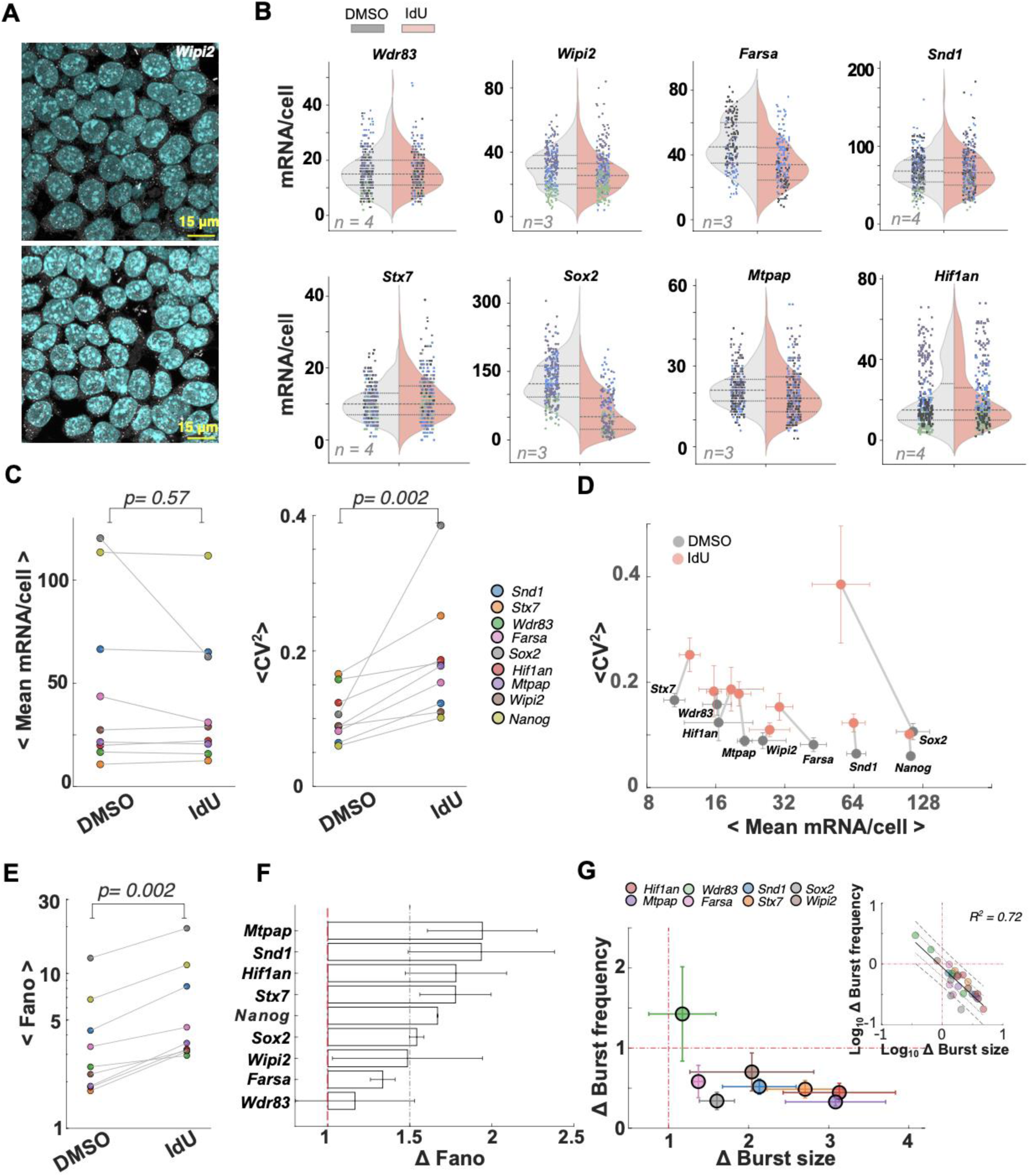
smFISH verification of noise amplification. **(A)** Representative smFISH images of *Wipi2* transcripts (white dots) in mESCs treated with DMSO (control, top) or IdU (treated, bottom), with DAPI stained nuclei (blue). Scale bar:15 μm. See also Fig. S4D. (See also Fig. S4) **(B)** Distribution of mRNA abundance per cell from all replicates in DMSO (grey) and IdU (red) treated samples. Colors in scatter correspond to replicates (n: number of replicates). Dashed lines represent median, first and third quartiles of mRNA distribution. **(C)** IdU induced changes in gene-expression quantified as mean mRNA/cell (μ) and increase in noise as mean CV^2^ (σ^2^/μ^2^); suggest statistical insignificance and significance respectively using the Paired Wilcoxon signed rank test on the means across replicates. Colors represent genes. **(D)** CV^2^-vs-mean mRNA abundance per cell for each gene is represented. DMSO (grey circles) and IdU (red circles). Error bars indicate ± SEM. **(E)** Increase in Fano factor (FF i.e., σ^2^/μ) in IdU treated cell-population is statistically significant from the untreated DMSO populations, inferred from paired, one-sided Wilcoxon signed rank test. (See also Fig. S5) **(F)** IdU/DMSO (i.e. fold change, as Δ) in Fano factor (i.e., FF IdU/FF DMSO) across the genes is plotted. Red-lines are guide to the eyes for axis of no-change. See also Fig. S5. **(G)** The fold change (Δ) in IdU to DMSO treated samples for burst frequency and burst size are plotted, as estimated from the fits to the negative binomial distribution. Error bars correspond to the SEM. Red-lines are guide to eyes for axis of no-change. (See also Fig. S6). *Inset:* The log-transformed fold-change (i.e., IdU/DMSO) in burst frequency and burst size are plotted for each gene from each replicate. Colors indicate genes. Solid black line indicates the fit to the linear regression with standard error of estimates (dashed and dotted lines for 2 and 1 respectively).

Despite previous observations that IdU-mediated noise amplification is largely independent of cell-cycle phase^20^, we nevertheless accounted for potential contributions from cell cycle. This analysis is based on an established body of literature that cell size is a surrogate for cell-cycle phase^11,20,39^ and we filtered out extrinsic noise by restricting the analysis to cells of similar size (**Fig. S4D**). At least three biological replicates were imaged to quantify per-cell mRNA abundance for each gene and condition with at least 50 cells used after cell-size based extrinsic noise filtering (**Fig. 2B).** This analysis revealed significantly greater variance and CV^2^ in mRNA levels in IdU-treated samples compared to controls for most genes while the change in mean-expression was insignificant (**Fig. 2C-D, Fig. S5A**). A direct comparison of mRNA CV^2^ values showed significant noise amplification for all the genes in the representative panel, without substantial changes in mean (**Fig. 2D, Fig. S5A**), in agreement with the scRNA-seq analysis. Notably, *Sox*2, and *Farsa* appeared to be exceptions exhibiting decreases in mean expression (see Discussion).

To ensure that noise amplification, as reported by CV^2^, could not be explained by changes in mean expression, we also analyzed the Fano factor calculated from the smFISH data (**Fig. 2E–F, S5B**). The absolute Fano factor was > 1 for all the genes, even in the DMSO-treated controls, indicating gene expression of the selected genes to be inherently bursty **(Fig. S5B)**, consistent with previous scRNA-seq and smFISH analyses of mESCs^20,40^. Moreover, Fano factor analysis verified that all genes in the panel exhibited significant IdU-induced amplification of noise (**Fig. 2E**). The greatest IdU-mediated noise amplification (i.e., fold-change or Δ in Fano factor) was for *Mtpap, Hif1an, Snd1, Stx7, Nanog* and *Sox2* (Δ Fano factor > 1.5) whereas *Farsa*, and *Wdr83* exhibited smaller noise amplifications **(Fig. 2F).**

To address, the possibility of over-estimation in the noise-metrics due to the variation in cell-size, even after filtering for extrinsic-noise, we computed corrected noise-metrics based on an established linear-regression analysis^39^. This analysis indicates no appreciable differences in the noise estimates (**Fig. S5C-D**), confirming lack of cell-size dependence after extrinsic noise-filtering, in contrast to a recent study^41^.

To further verify IdU-mediated noise amplification, we tested if the mechanistic underpinnings^19^ of homeostatic noise amplification were satisfied. Theory predicts that homeostatic noise amplification (i.e., changing noise without altering mean level) requires reciprocal changes in at least two transcriptional bursting parameters^19^; e.g., a decrease in transcriptional burst frequency coupled with a corresponding but reciprocal increase in transcriptional burst size such that the mean number of transcripts remains unchanged. Specifically, the two-state random telegraph model could fit mESC smFISH data but to account for IdU-mediated noise amplification, inclusion of an additional “off” state in the model coupled with a feedforward gain was required^20^. Here, based on established literature^6,40^, we first inferred the effective burst size and frequency by fitting the mRNA distributions to a negative binomial distribution (**Fig. 2G**, **Fig. S6A,C**) and then quantified the relative IdU-mediated change in noise—importantly, the negative binomial fitting approach does not explicitly account for underlying molecular mechanisms such as the additional off state or feedforward gain. The fitting further shows a reciprocal relationship between the fold change in burst size and frequency **(Fig. 2G)** in agreement with the model of homeostatic noise amplification. The one exception was *Wdr83* which showed a different pattern of burst size and frequency and is discussed below.

Overall these smFISH data further validate the scRNA-seq analysis of transcriptional noise and support that IdU generates a highly penetrant homeostatic amplification of transcriptional noise for 8 out of 9 genes in this panel of genes selected from diverse expression profiles and locations in the genome.

## DISCUSSION

This study set out to determine: (i) generally, which scRNA-seq algorithm was most appropriate for quantifying transcriptional noise using the noise-amplifying molecule IdU^20^ as a perturbation; and (ii) specifically, if the IdU noise-enhancer molecule acts in a globally penetrant manner.

Analysis of scRNA-seq data using different algorithms indicated that the amplification of noise by IdU is globally penetrant and likely not an artifact of a particular scRNA-seq analysis algorithm. However, the scRNA-seq analysis also indicated that scRNA-seq algorithms generate variable genome-wide noise profiles (**Fig. 1**), which each algorithm suggesting a quantitatively different % penetrance. A second scRNA-seq analysis in Jurkat cells (**Fig. S3**) was consistent with IdU acting as a global noise enhancer, independent of the specific biological context. To validate the scRNA-seq noise analysis, we next used smFISH to examine a panel of genes that exhibited high sensitivity to individual scRNA-seq algorithms and represented both high and low expressing genes. smFISH analysis revealed that IdU increased transcriptional noise for 8 out of 9 genes (**Fig. 2**). Overall, the results indicate that most published scRNA-seq algorithms were fairly accurate for analysis of gene-expression noise compared to the smFISH direct measurement (**Fig. 3A-B**). Notably, the estimates from the “*raw* method” are similar to those from the other algorithms, although all the methods tend to systematically underestimate the fold-change in noise metrics compared to smFISH. However, a combined score based on the minimal deviation from the smFISH estimates suggest that “*raw”* and “*BASiCS”* perform relatively better among all the other algorithms tested **(Fig. 3C)**. In BASiCS, the fold-change in mean, CV^2^ and Fano were directly estimated from normalized counts (using the normalization parameter obtained from the Markov Chain Monte Carlo (MCMC) fits), and resulted in minimal data-transformation (**Fig. S1**). Additionally, MCMC fits provide robust estimates for all the model parameters such as the overdispersion (i.e., gene-specific parameter for biological noise that accounts for noise beyond the Poissonian noise after accounting for technical noise, analogous to CV^2^), that indicates ∼84 % of the genes exhibit a fold-increase in biological noise upon IdU perturbation, which is comparable to the direct-CV^2^ (as in **Fig 1A**, ∼86 %). Further, the variance decomposition indicates that ∼26 % and ∼75 % of the total genes show IdU-induced amplification for Poissonian and super-Poissonian components, respectively, while only a ∼3 % exhibited an increase in both. Overall, this study suggests that direct estimates of noise along with the BASiCS metrics such as overdispersion, residual variability, and variance decomposition, provide accurate quantification of transcriptional noise and insights into the phenomenology underlying the gene expression variability.

**Figure 3:**
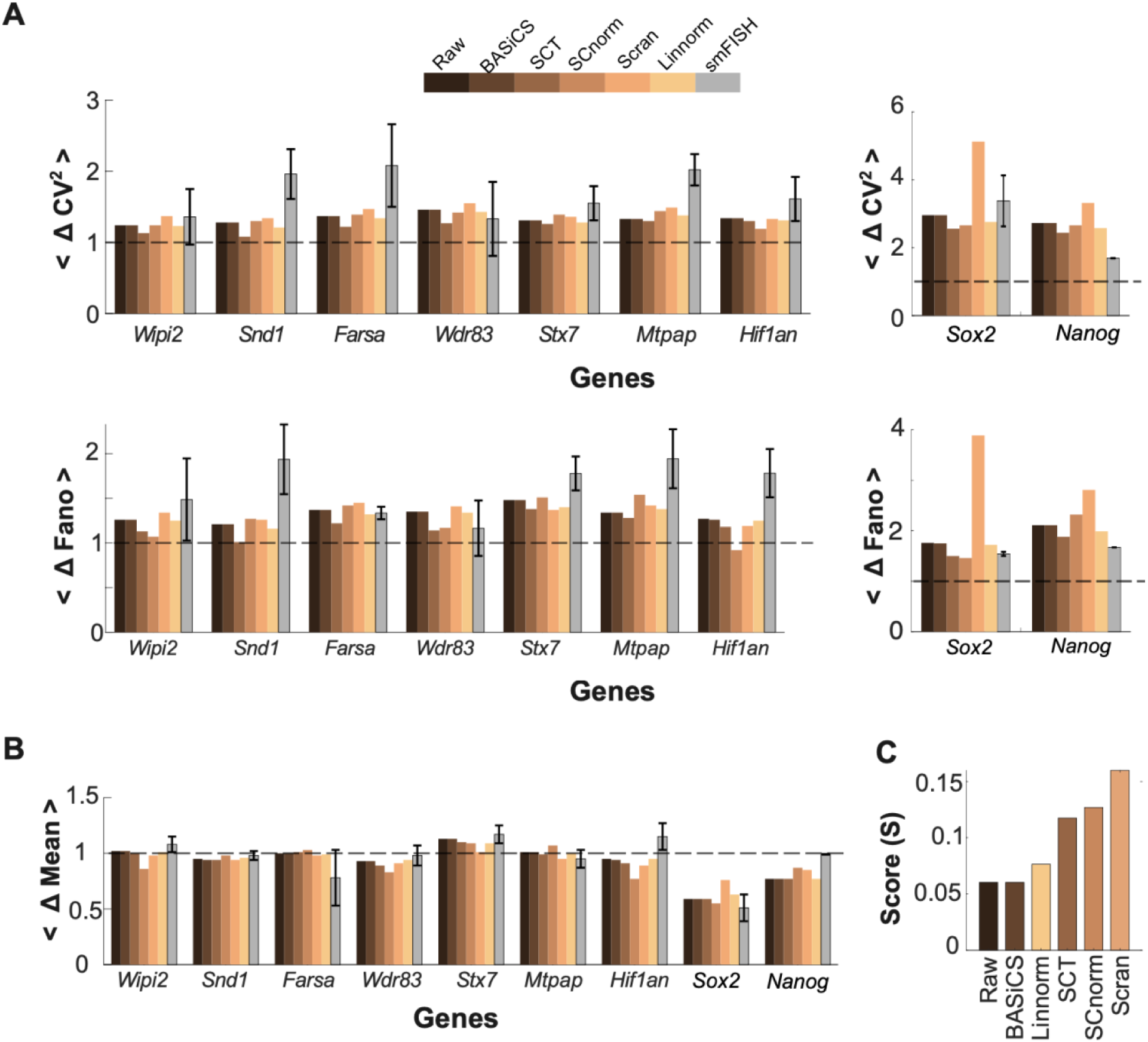
Comparison of noise amplification between smFISH and scRNAseq. The fold-change (Δ) in **(A)** noise metrics and **(B)** mean obtained from different scRNA-seq algorithms is compared to the estimates from measured smFISH across the panel of selected genes. **(C)** Bar plot depicts the score, (*S*) comparing the performance of each scRNA-seq method to the FISH based on the deviation for the three metrics (described in methods section).

The Jurkat scRNA-seq analysis of noise (**Fig. S3D**) is generally consistent with previous smFISH analysis in Jurkats indicating that 75% of genetic loci have a Fano factor > 1^11^ and the two scRNA-seq biological replicates of the Jurkat experiment confirm that the noise difference between IdU and DMSO samples cannot be attributed to technical bias of the single mESC DMSO and IdU samples alone. However, Jurkats appear to exhibit a significantly lower extent of noise amplification. Several mechanisms may account for the observed difference. First, slowly dividing cells, such as Jurkats, have slower DNA replication rates, thus IdU incorporation has less impact on transcriptional dynamics due to the relatively fast activity of the base-excision repair (BER) pathway through which IdU is removed from the genome. Second, a technical difference between the DMSO and IdU-treated mESC cells is sequencing depth—while the IdU-treated mESCs were sequenced more deeply, they exhibit lower median UMI count per cell compared to the DMSO-treated cells (**Fig. S3E**), suggesting a possible reduction in cellular total RNA content upon IdU treatment. This reduction suggests the possibility of lower absolute mean-RNA levels masked by normalization methods, and higher noise stemming from Poisson scaling, which inflates the noise enhancement effect in the mESC dataset. Third, another technical limitation of our Jurkat scRNA-seq data is the moderate sequencing depth (**Fig. S3E**), allowing noise quantification in only ∼1000 genes, compared to the ∼4000 genes in mESCs. These ∼1000 genes comprise the highest expression percentile of all expressed genes. It is possible that noise amplification is less efficient for these very abundant transcripts due to higher RNA stability and/or more continuous than bursty transcription^42^. Finally, there is also a possibility that noise amplification is mESCs could potentially propagate into cell-state alterations in mESCs, which may introduce transcriptional heterogeneity in these pluripotent stem cells and could reflect potential biological changes in some cells^43^ (a possibility that may be less likely to occur in terminally differentiated Jurkat cells). However, we feel this possibility is unlikely since these biological changes in mESCs were not previously observed^20^ and would likely be reflected by changes in mean-expression levels which were not observed here or previously^20^.

The smFISH replicates, performed on two different microscopes each using different confocal technologies (i.e., spinning disk versus laser scanning), generated similar estimates in fold-change of noise across the panel of genes, which indicates robustness and lack of bias that emerge from the acquisition differences. The smFISH analysis also allowed the calculation of fold change in burst-size and burst frequency for selected genes (**Fig. 2**) and the results were qualitatively comparable to the estimates from our scRNA-seq in an independent recent study using mechanistic models for estimation of the bursting kinetics^44^ with a quantitative match for the intermediate-abundance gene *Mtpap* (i.e., Log_2_ fold change in burst size and frequency in smFISH: 1.5 and - 1.6; while the estimates from Monod pipeline are 1.3 and −1.5 respectively). The smFISH data verify that IdU amplifies intrinsic transcriptional noise homeostatically (i.e., without altering mean expression level) for most genes irrespective of genomic location and expression level.

One technical limitation of this study is that smFISH analysis is necessarily low throughput and limited to the subset of genes for which good probe sets can be designed, which limits the spectrum of measurements and the number of genes that can be analyzed. We attempted to mitigate this by exploring genes from across the expression spectrum (**Fig. S4A–B**), and by analyzing a substantial number of cells per treatment (i.e., at least ∼50 cells per condition and replicate after filtering extrinsic noise) and the estimates from replicates are consistent with statistics obtained from pooled datasets (**Fig**. **S6E**). Thus, the data herein indicate that IdU likely acts as a noise-enhancer molecule for a large fraction of genes irrespective of their mean-expression level.

However, the degree of noise amplification varies. A higher fold-change in Fano is observed for *Mtpap and Hif1an* genes associated with stress and metabolic processes*, Syndapin and Syntaxin7* involved in the endocytosis and vesicular trafficking respectively. The pluripotency associated transcription factors such as *Sox2* and *Nanog,* exhibit intermediate noise-amplification. While, the genes associated with translation and autophagy i.e., *Farsa* and *Wipi2*, and *Wdr83* which is involved in mRNA-processing and functions as a molecular scaffold - exhibit lower-noise amplification. Interestingly, magnitude of protein noise in yeast cells is higher for the stress and metabolism associated proteins and lower for those involved in protein complexes and translation machinery^13,14^, suggesting a potential role of IdU perturbation in investigating the contribution of transcriptional and translational noise. Notably, *Wdr83* is an outlier for its change in noise (**Fig. 2F**) and the estimates of the change in burst size and frequency do fall outside the reciprocal change expected for orthogonal noise amplification (**Fig. 2G**). Considering that IdU increases noise via BER, a genome-wide surveillance pathway, it may be surprising that any gene fails to exhibit an increase in noise. It is possible that the low-throughput nature of smFISH may have obscured IdU-induced noise amplification or the variation could be inherent to the gene. When replicates for each gene were pooled, the homeostatic increase in noise was maintained (i.e., increase in both Fano factor and CV^2^ without a significant change in mean) for all genes except *Wdr83,* and the reciprocal relationship between burst size and frequency was also maintained except *Wdr83* (**Fig**. **S6E**). However, it should be noted that these population statistics include 200–300 cells per gene/condition and do suffer from some inter-replicate variability. A potential explanation for the *Wdr83* outlier may lay in mechanism by which BER increases noise via DNA topology changes. Specifically, the BER enzyme AP endonuclease 1 (Apex1) generates DNA supercoiling which leads to an accumulation of RNA Polymerase II; when released, this amplifies transcriptional burst size; indeed, the IdU noise effect can be phenocopied by topoisomerase inhibitors, which increase DNA supercoiling^20^. Consequently, regions which naturally have increased supercoiling and corresponding high levels of topoisomerase may be less sensitive to changes in topology caused by IdU and BER-induced supercoiling. It has been previously reported that Topologically Associated Domain (TAD) boundaries are regions that exhibit substantial supercoiling and are enriched in insulator binding protein CTCF^45,46^, and that Topoisomerase IIB is prevalent at CTCF sites^47^. Intriguingly, *Wdr83* has two relatively unique features among the panel of genes tested: (i) it contains a relatively large CTCF binding region, and (ii) it is found at a TAD boundary^45^. Together these two features may be consistent with IdU acting as a topology-dependent global noise-enhancer molecule, as shown by scRNA-seq **(Fig. 1)** and smFISH **(Fig. 2)** analysis.

A second point of interest is the case of *Sox2.* Whilst most genes analyzed exhibited a homeostatic increase in noise—without a substantial change in mean—smFISH revealed that IdU induced a decrease in mean *Sox2* mRNA, though we note the change in mRNA numbers was less than two-fold. Notably, IdU did not alter single-cell *Sox2* protein levels in our previous analysis^20^. It is possible that the reported negative-feedback regulation of *Sox2*^48^ acts to buffer changes at the protein level.

From the practical perspective of quantifying expression noise, this study reveals that common analyses can fail to resolve quantitative changes in intrinsic expression noise, particularly for individual genes. A number of approaches have been proposed to overcome these types of scRNA-seq limitations, including elegant solutions using mathematical modeling methods to address technical variability without compromising quantification of biological noise^40^, though these can be cumbersome to implement. The analyses herein indicate that it may be advisable to combine high-throughput analyses (e.g., scRNA-seq) with lower-throughput direct quantification (e.g., smFISH) to quantify changes in transcriptional noise for individual genes as the “ground-truth” may lie somewhere in between the results of each analysis. Regardless, both the scRNA-seq and smFISH data argue that IdU appears to be a global noise enhancer which could be leveraged to modulate noise without altering mean expression for the majority of genes.

### Limitations of the study

While all the scRNA-seq algorithms explored in this study indicate IdU results in genome-wide homeostatic amplification of transcriptional noise, it is limited by a single replicate in mESCs and smaller sequencing depth in the two replicates of Jurkat. Additionally, although smFISH estimates support the scRNA-seq results in mESCs, this validation is restricted to a smaller set of target genes due to technical limitations, which may also constrain the evaluation of the algorithms using smFISH as a reference. Furthermore, variations observed across the replicates in the *Wdr83* gene along with its contrary trend from the scRNA-seq estimates raise the questions on the contribution of topoisomerase-based relaxation of super-coiling upon IdU-induced repair versus the accessibility and effectiveness of IdU itself at the TAD regions, as well as the influence of the biophysical properties of the gene. Our analysis is limited to the relative estimates of burst size and frequency, inferred from the fitting of mRNA distributions to a Negative Binomial distribution under the assumption that mRNA stability is unaltered upon IdU-treatment, due to the lack of experimentally measured half-life estimates for all the target mRNAs. Lastly, while our results indicate a global penetrance of IdU, it is based on the fast-dividing embryonic mouse stem cells and transformed human Jurkat cells. The extent of IdU penetrance and quantification of transcriptional noise in primary differentiated cell lines, which usually have relatively slower proliferation rate, shorter replication dwell times, and different chromatin landscapes and dynamics, remains to be explored for utilizing IdU as a noise-enhancer probe.

## Acknowledgements

We thank Ravi Desai, Gustavo Vasen, Karla Delucas, and Daniel Lewis for technical guidance. Kathryn Claiborn for editing and Oded Regev for useful discussions. Data for this study were acquired at the Nikon Imaging Center at UCSF using NIH S10 Shared Instrumentation grant (1S10OD017993-01A1). Gladstone Genomics Core performed 10x Genomics library construction. Sequencing was performed at the UCSF CAT, supported by UCSF PBBR, RRP IMIA, and NIH 1S10OD028511-01 grants.

## Funding

This work was supported by NIH award R37AI109593 (to LSW), BZ acknowledges support from the EMBO fellowship (ALTF 388-2021) and XC acknowledges support from CIRM training grant EDUC4-12766. XGA acknowledges support from NIH training grant K12GM081266.

## Author contributions

GC, XC, and LSW conceived and designed the study. GC and XC designed the smFISH experiments. GC and NK performed, analyzed, and curated the data. GC and BZ performed mESCs scRNA-seq normalizations and curated the data. BZ performed Jurkat scRNA-seq experiments, analyzed, and curated the data. LSW provided reagents and resources. GC, XC, NK, BZ, XG and LSW wrote the paper.

## Declaration of Interests

The authors declare that they have no competing interests. L.S.W is an equity co-founder of VxBiosciences Inc and Autonomous Therapeutics Inc.

## Supplemental information

Document S1. Figures S1–S6 Dataset S1-S3.

## STAR METHODS

| REAGENT or RESOURCE | SOURCE | IDENTIFIER |
| --- | --- | --- |
| Critical commercial assays |  |  |
| Chromium Next GEM Single Cell 3' HT Kit v3.1 | 10x Genomics | 1000370 |
| NovaSeqX 10B flow cell (v1.5 reagents) | Illumina | 20085594 |
| 10x Cell Ranger (6.1.2, 7.0.1) | 10x Genomics | https://www.10xgenomics.com/support/software/cell-ranger/downloads#download-links |
| Chemicals, peptides, and recombinant proteins |  |  |
| DAPI | Thermo Fisher Scientific | Cat#D1306 |
| IdU | Millipore Sigma | Cat# I7125 |
| Formaldehyde | Tousimis | Cat#1008A |
| Formamide (deionized) | Ambion | Cat#AM9342 |
| 20x SSC buffer | Invitrogen | Cat#AM9770 |
| Glycerol | Thermo Fisher Scientific | Cat#17904 |
| Glucose oxidase | Sigma Aldrich | Cat#G7141 |
| Catalase | Sigma Aldrich | Cat#C3515 |
| Imaging chamber (8-well dish) | Ibidi | Cat#80826 |
| Deposited data |  |  |
| Single-cell RNA-seq Jurkat | This study | GSE263194 |
| Single-cell RNA-seq mESC | Desai et al. 2021<sup>20</sup> | GSE176044 |
| Experimental models: Cell lines |  |  |
| mESC | Desai et al. 2021<sup>20</sup> |  |
| Jurkat | American Type Culture Collection | TIB-152 |
| Oligonucleotides |  |  |
| smRNA-FISH probes, See Data S3 | This study ( Stellaris, LGC Biosearch Technologies) | N/A |
| Software and algorithms |  |  |
| Fiji/ImageJ | Schneider et. al. 2012<sup>49</sup> | https://imagej.nih.gov/ij/ |
| MATLAB (R2024a) | MathWorks<sup>50</sup> | https://www.mathworks.com/products/matlab.html |
| R (4.4.0) |  |  |
| FISH-Quant ( version – v2a ) | Mueller et. al. 2013<sup>38</sup> | https://bitbucket.org/muellerflorian/fish_quant/src/master/ |
| BASiCS (version 2.15.5) | Vallejos et. al. 2015<sup>31</sup>, Eling et. al. 2019<sup>32</sup> | https://www.bioconductor.org/packages/3.7/bioc/html/BASiCS.html |
| Linnorm (2.28.0) | Yip et. al. 2017<sup>30</sup> | https://www.bioconductor.org/packages/release/bioc/html/Linnorm.html |
| SCnorm (1.26.0) | Bacher et. al. 2017<sup>33</sup> | https://www.bioconductor.org/packages/release/bioc/html/SCnorm.html |
| SCTransform (0.4.1) | Hafemeister, C., and Satija, R, 2019<sup>28</sup> | https://cran.rproject.org/web/packages/sctransform/index.html |
| Scran (1.30.2) | Lun et. al. 2016<sup>29</sup> | https://bioconductor.org/packages/release/bioc/html/scran.html |

## Resource availability

### Lead contact

Further information and requests for resources and reagents should be directed to and will be fulfilled by the lead contact, Leor Weinberger.

### Materials availability

This study did not generate new unique reagents

### Data and code availability

Jurkat single-cell RNA-seq raw and processed sequencing data have been deposited at GEO and are publicly available as of the date of publication. Accession numbers are listed in the key resources table. Microscopy data reported in this paper will be shared by the lead contact upon request. Any additional information required to reanalyze the data reported in this paper is available from the lead contact upon request.

## Method details

### Cell culture

Mouse embryonic stem cells (E14, male) were cultured on gelatin-coated platers with ESGRO-2i/LIF medium (Millipore, cat: SF002-500) at 37°C, 5% CO2, in humidified conditions, as described previously^20^. Jurkat T Lymphocytes were cultured in RPMI-1640 medium (supplemented with L-glutamine, 10% fetal bovine serum, and 1% penicillin-streptomycin), at 37°C, 5% CO_2_, in humidified conditions. Cells were treated with either DMSO or 10/20 μM IdU (I7125, Millipore Sigma) for 24/48h in mESC/Jurkat respectively.

### Single-Cell RNA Sequencing Analysis (mESCs and Jurkats)

For mESCs, we used our previously published^20^ deeply sequenced scRNA-seq dataset available at the GEO repository under accession number GSE176044. Read mapping to mm10 genome was done with CellRanger (6.1.2)^51^ to obtain a gene count matrix. Before applying each normalization method, both DMSO and IdU datasets were subjected to a quality control process. First, Seurat^52^ was used to filter for high-quality cells using a minimum of 4000 detected genes, 10000 UMI counts, and <10% reads mapping to mitochondrial genes per cell. The resulting count matrix was then further filtered by the ‘BASiCS_Filter’ function from the BASiCS^31^ R package with default parameters, which limited the analysis to genes with sufficient sequencing coverage for reliable noise quantification. The output count matrix consisted of 811 cells for DMSO and 732 cells for IdU across 4456 genes. This output was then used to run five normalization pipelines according to their protocols: SCTransform^28^, scran^29^, Linnorm^30^, BASiCS^31^, and SCnorm^33^. For comparison, the filtered output was also normalized using a simple approach referred to as the “raw” method, here. In this method, the gene-specific counts in each cell were divided by the total counts in that cell and then scaled by a factor of 10^4^. BASiCS was run with default parameters and recommended settings (N=20000, Thin=20 and Burn=10000) using the horizontal integration strategy (no-spikes). The normalized data obtained from ‘BASiCS_DenoisedCounts’ function was further normalized similar to the raw data. The other packages were run with default parameters.

For Jurkats, two biological replicates (4 samples) of single-cell suspensions were loaded on a Chromium X instrument using a Chromium Next GEM Single Cell 3’ HT Kit v3.1 (10x Genomics). Sequencing was performed on an Illumina NovaSeq 6000 with a paired-end setup specific for 10x libraries. Data were aligned to hg38 reference genome using 10x Cell Ranger (7.0.1). Resulting sequencing depth as estimated by median UMI count per sample is shown in Fig. S3E. Analysis was performed similarly to the mESC dataset: Seurat was used to filter for high-quality homogeneous cell population using the following filters: 3000 < detected genes < 5000, 8000 < UMI counts < 15000, and <10% reads mapping to mitochondrial genes per cell. The resulting count matrix was then further filtered by the BASiCS_Filter function with default parameters. The output count matrix consisted of 5988 and 5786 cells for DMSO samples and 5148 and 4701 cells for IdU samples across 1107 genes. This output was then used to run the above-mentioned 5 normalization pipelines according to their protocols. For each sample, we calculated separately for each biological replicate the mean, CV^2^, and Fano values per gene and then averaged the values of both replicates for further analysis.

### Single Molecule RNA FISH

Cells within 3 to 12 passages were used for smRNA FISH experiments. Probes for the detection of transcripts were developed using the designer tool from Stellaris (LGC Biosearch Technologies) (Data S3) setting the minimum number of probes to 30 (TAMRA conjugated) for gene transcripts. 1.5 x10^5^ mouse embryonic stem cells were seeded into well of a gelatin-coated, 8-well Ibidi dish (cat: 80826) in 2i/LIF media. 24 hours following seeding, media was replaced with 2i/LIF containing 10 mM IdU or equivalent volume DMSO. After 24 hours of treatment, cells were then fixed with DPBS in 4% paraformaldehyde for 10 minutes. Fixed cells were washed with DPBS and stored in 70% EtOH at 4°C for one hour to permeabilize the cell membranes. Probes were diluted 200-fold and allowed to hybridize at 37°C overnight. Wash steps and DAPI (Thermo, cat: D1306) Wash steps and DAPI (Thermo, cat: D1306) staining were performed as described (https://www.biosearchtech.com/support/resources/stellaris-protocols). Briefly, after 16 hours, cells were washed with wash buffer and incubated for 30 minutes at 37°C twice, followed by DAPI stain (DAPI in wash buffer at 10 ug/ml), for 15 minutes at 37°C. The cells were washed with 2x SSC (Invitrogen, cat: AM9770) once, followed by freshly prepared GLOX solution and incubated for 3 minutes. The cells were finally suspended in the anti-fade GLOX buffer with enzymes (i.e., also the imaging buffer) to minimize photo-bleaching (buffer containing 50% glycerol (Thermo, cat: 17904), 75 mg/mL glucose oxidase (Sigma Aldrich, cat: G7141), and 520 mg/mL catalase (Sigma Aldrich, cat: C3515). Images were collected on an inverted Nikon TiE microscope (Nikon) run using Micromanager 2.0^53^ equipped with a CSU-W1 Spinning Disk with Borealis Upgrade (Yokogawa, Andor), ILE Laser launch with 4 laser lines (450/488/561/646nm, Andor), quad-band dichroic ZT405/488/561/647 (Chroma), emission filters for DAPI (ET447/60), GFP (ET525/50), RFP (ET607/36), and Cy5 (ET685/40) (Chroma), piezo XYZ stage (ASI), and Zyla 4.2 CMOS camera (Andor), using a Plan Apo VC 60x/1.4 Oil objective (Nikon). Approximately 10 XY regions of interest were randomly selected for each condition. For each image, XY pixel size was 108nm/px, and a Z-step size of 250nm was used with over 60 image planes to fully cover the tissue. The additional replicates were imaged on a confocal laser scanning microscope (Fluoview 3000 Olympus™) with a 63x (1.4 NA) oil-immersion objective. Approximately, 10-15 XY regions were randomly selected for each condition at z-step size of 280nm for about 40 – 60 frames to span the cells. Prior to imaging, cells were checked for Nanog-GFP presence to confirm the pluripotent state of the mESCs.

### Image analysis and extrinsic noise filtering for quantification

Image analysis and spot counting was performed using FISH-quant^38^. Images, were background subtracted with a rolling ball radius of 10 pixel in the TAMRA channel for the control and treated sets requiring pre-processing in Fiji^49^. Cells were manually segmented to ensure selected cells for the analysis were: (i) single and non-overlapping in all dimensions; (ii) non-dividing (based on cell shape) and those at the later stages of cell-division (such as metaphase and anaphase) were excluded based on DAPI staining; and (iii) of similar size to minimize extrinsic noise. To remove outliers, for every pair of probes per replicate, cells with areas below 5^th^ and above 95^th^ percentile (calculated from the combined DMSO and IdU population) were excluded. This was followed by iteratively eliminating cells with 2 and 98 percentile thresholds until the area distributions of DMSO and IdU satisfied statistical insignificance (Permutation test, a = 0.01, number of permutations with replacement = 10000) and the correlation coefficient for the linear relationship between mRNA abundance and cell-size was less than < 0.45 (24/28 cases had P > 0.05 and R^2^ < 0.3). This approach does not assume a prior-distribution and removes any bias that could stem from differences in the number of cells per group. For, all the analysis, gene-sets with at least >50 cells/treatment were considered, (except for one *Farsa* and *Syntaxin7* replicate with ∼ 25 and ∼ 35 cells/condition). The lack of cell-size dependent effects was further confirmed by computing the cell-size corrected noise-metric as described^11,39^, which yielded no changes in the measured metrics per gene per replicate. (a total of 2201 DMSO and 2035 IdU treated cells were analyzed after extrinsic filtering from a total of 2748 and 2693 segmented cells.)

### Estimation of Burst size and Burst frequency

The distribution of mRNA/cell for each gene and replicate were fit to the negative binomial distribution using maximum likelihood estimates (nbinfit, MATLAB). The estimated burst size and burst frequency from the fits were inferred as: mean = burst size x burst frequency and Fano factor is 1 + burst size (Fig. S6A,C). Data from all replicates was also pooled for each gene and the mRNA distributions fit to estimate the bursting-parameters.

### Quantification and Statistical Analysis

Statistical analysis was performed in MATLAB. To test for the homeostatic noise amplification, increase in transcriptional noise between DMSO and IdU treated samples in smFISH data, the mean of the measures over the replicates were considered. A non-parametric paired Wilcoxon signed rank was used to test for significance in mean (two-sided). While a one-sided paired Wilcoxon signed rank test was used to test for the significance for increase in noise and burst size, and decrease in burst-frequency (based on the mechanism of IdU shown previously^20^ and from the scRNA-seq analysis in this study). The Permutation test was used to filter extrinsic noise and test for the significance in cell-area distributions between DMSO and IdU cells in smFISH data. To evaluate the performance of the scRNA-seq methods for homeostatic noise amplification, a combined score (S) was computed for each method (k); defined as the linear sum of the medians 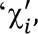 computed for each metric (i.e., *‘i’*). The metrics 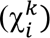 are relative deviations in mean expression (μ), squared coefficient of variation 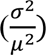 and Fano factor 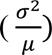 from scRNAseq method 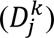 and smFISH (*E*_*j*_) across all the genes (*‘j’*). The minimal score *S*_*k*_ corresponds to the method that matches the smFISH closely.

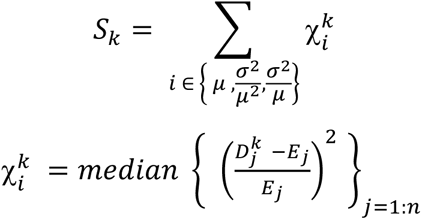

## Notes

### Competing Interest Statement

The authors have declared no competing interest.

