## Supplementary Material for "Quantitative comparison of single-cell RNA sequencing versus single-molecule RNA imaging for quantifying transcriptional noise"

### SUPPLEMENTAL INFORMATION

- Supplemental Figure Legends

- Supplemental Figures

- Supplemental Datasets

### SUPPLEMENTAL FIGURE LEGENDS

**Figure S1: Comparison of scRNA-seq algorithms for noise quantification**. Each panel shows a comparison between fold changes in $CV^2$ (upper-right panels) and Fano (lower-left panels) in response to IdU treatment as quantified by the indicated methods. The dashed green lines indicate the raw-data axes for reference. Related to Figure 1E.

**Figure S2: Reproducibility of noise quantification metrics._(A-B)** Randomly sampled cells from the mESC DMSO sample show no significant changes in $CV^2$ **(A)** or Fano **(B)** values between the two subpopulations. **(C-D)** Two biological replicates of DMSO-treated Jurkat cells allow validation of the reproducibility of noise quantification using $CV^2$ **(C)** or Fano **(D)** as noise metrics. **(E)** Sequencing depth as estimated by median UMI counts of the Jurkat and mESC samples used in this study. Values correspond to initial read counting as summarized by CellRanger (see Methods section for details).

**Figure S3: scRNA-seq analysis in Jurkat cells.** **(A)** scRNA-seq analysis of $CV^2$-vs-mean for ~1,000 transcripts in Jurkat cells treated with IdU (red) or DMSO control (black) as analyzed by commonly used normalization algorithms: SCTransform, scran, Linnorm, BASiCS, or SCnorm. **(B)** Mean expression for each of the ~1,000 transcripts in presence and absence of IdU using each normalization algorithm; none of the normalization algorithms generate significant changes in mean expression for IdU-treated cells. **(C)** $CV^2$ for each transcript in presence and absence of IdU using each normalization algorithm; different algorithms generate slightly different fractions of transcripts with amplified noise ranging from ~59% of transcripts with amplified noise (scran) to ~66% of transcripts with amplified noise (SCnorm). Wilcoxon rank sum tests comparing the DMSO and IdU $CV^2$ values for all genes are of marginal statistical significance ($P < 0.005$ for SCT, BASiCS and SCnorm; $P > 0.01$ for Linnorm and scran). **(D)** Quantification showing percentages of transcripts with indicated Fano factor fold changes between IdU-treated and control cells. Wilcoxon rank sum tests comparing the DMSO and IdU Fano values for all genes are highly significant for all algorithms ($P < 10^{-17}$) besides scran ($P=0.013$). **(E)** Pearson correlation coefficients between fold changes in noise metrics between IdU-treated and control cells. Shown are correlations between the indicated methods quantifying fold changes in $CV^2$ (left heatmap) and Fano (right heatmap).

**Figure S4: Representative smFISH images for selected panel of genes and extrinsic noise filtering. (A)** IdU-induced amplification of noise for selected panel-of 8 genes as quantified by Fano factor of BASiCS analysis of mESC data with their genomic location for smFISH validation; Nanog data is from Desai et. al., (2021). **(B)** Direct comparison of fold-change in mean and Fano factor from five scRNA-seq algorithms for the eight genes for smFISH validation; Nanog data is from Desai et. al., (2021). **(C)** Maximum intensity projections of representative smFISH images of mESCs treated with DMSO and IdU ( top and bottom) for 24 hours with DAPI stain (blue) and mRNA transcripts (grey) labeled with TAMRA probe-set. are shown. Scale bar: 15 μm **(D)** Extrinsic noise filtering based on cell-area distributions for *Wipi2* from a replicate is represented (pre- filter: DMSO:178 cells, IdU: 179 cells, $p < 10^{-4}$ from permutation test and after extrinsic noise filtering: DMSO:108 cells, IdU: 105 cells, $p = 0.11$ ). Related to Figure 2.

**Figure S5. IdU mediated orthogonal increase in noise across the selected panel of genes**. **(A)** mean-mRNA per cell as by smFISH between DMSO and IdU treated populations for selected genes. Colors represent genes. **(B)** Fano factor versus mean mRNA abundance per cell for the selected genes. All the error bars correspond to SEM. **(C)** IdU-induced change in change in $CV^2$ (left) and Fano factor (right). Error bars correspond to SEM. **(D)** Cell-size corrected (i.e., extrinsic noise filtered) change in $CV^2$ (left) and Fano factor (right). Error bars correspond to SEM. Related to Figure 2.

**Figure S6. Reciprocal changes in burst size and frequency upon IdU treatment from smRNA FISH estimates. (A-B)** Burst size and **(C-D)** Burst frequency estimated from negative binomial fitting to the mRNA distributions for DMSO (grey) and IdU (orange) treated cells. Bars correspond to the mean across replicates. Error bars represent SEM. The increase in burst size and decrease in burst frequency are statistically significant using one-sided paired Wilcoxon signed rank test. **(E)** i.e. mRNA/cell across replicates for each gene (left). Red lines represent axis of no-change. Right: fold-change in mean, $CV^2$ and Fano factor from the pooled data-set. Related to Figure 2.

### SUPPLEMENTAL DATASETS

**Data S1. Processed data – mESC.** Statistics of gene expression counts normalized by the indicated method in the "method" column. Columns contain mean, $CV^2$ and Fano values for DMSO and IdU samples, as well as the log2 fold-changes (IdU/DMSO) for each of the statistics. "diff_Fano_2" and "diff_Fano_bin" contain fold-changes (not log2) used for Fig. 1D.

**Data S2. Processed data – Jurkat.** As Data S1, for Jurkat dataset. Associated with Fig S2D.

**Data S3. smRNA-FISH Probe List.** Sequences of smRNA-FISH oligonucleotide probe for eight genes

### Supplemental Figures

#### Figure S1

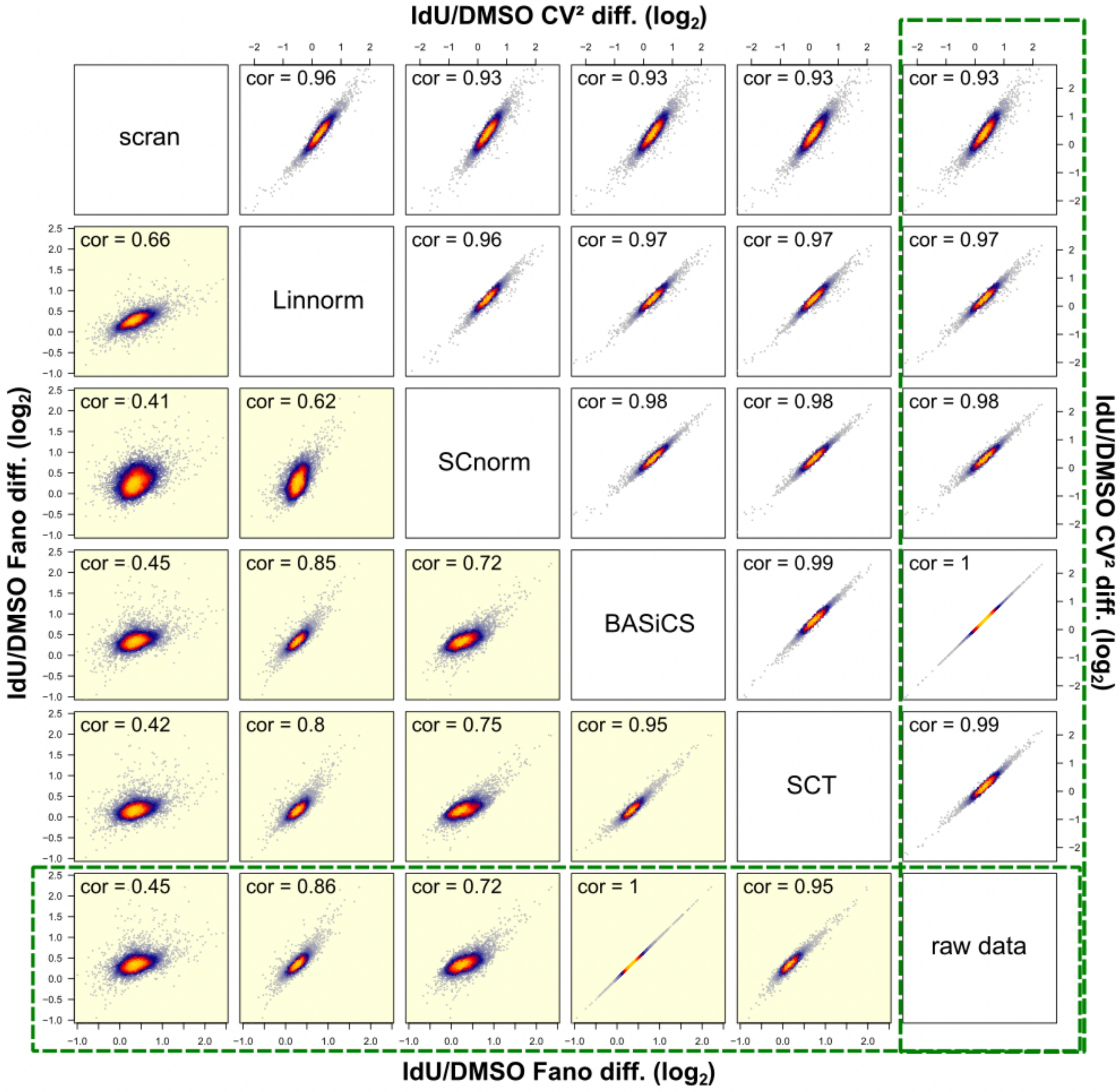

**Figure S2**

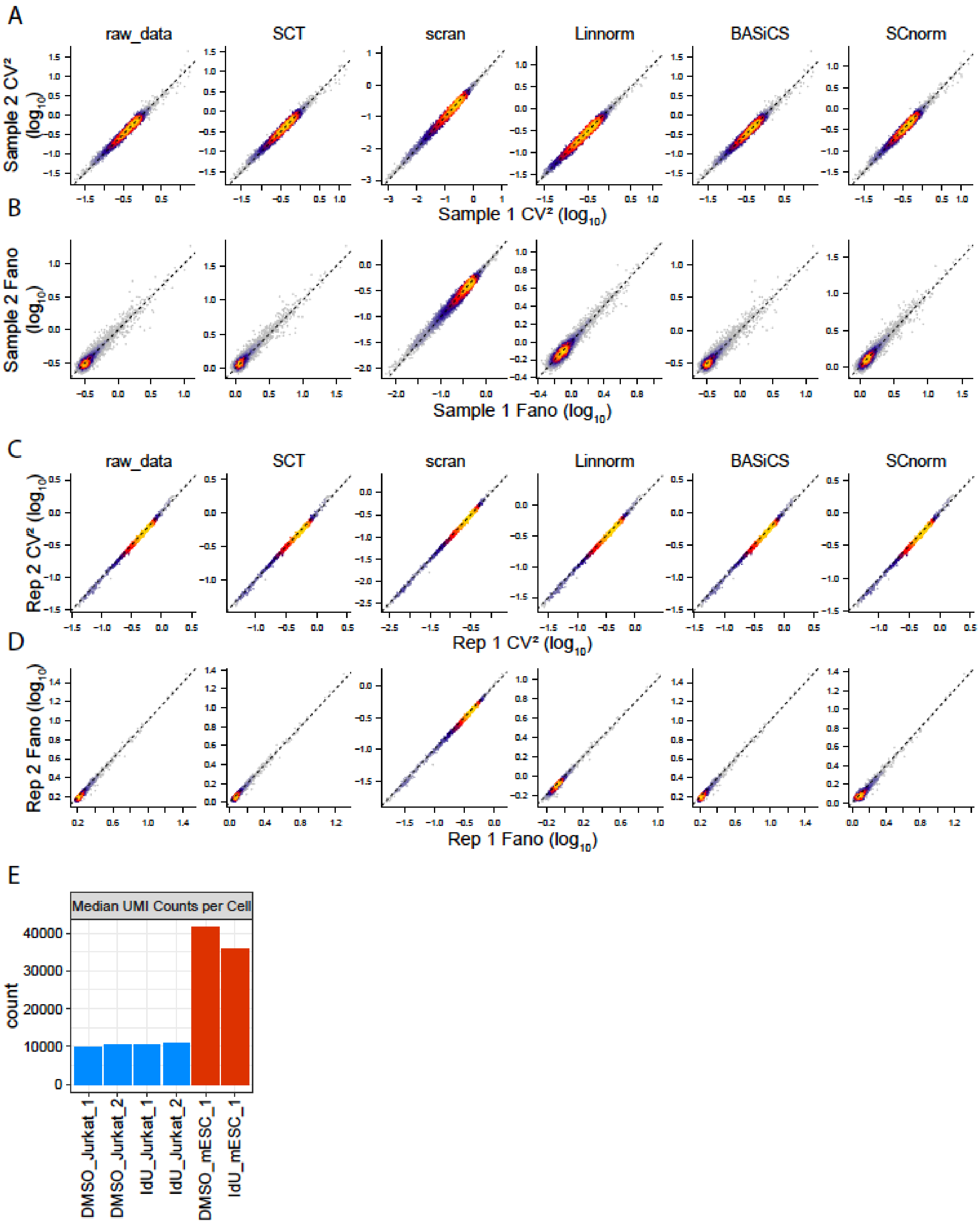

A
raw_data
SCT
scran
Linnorm
BASiCS
SCnorm
Sample 2 CV² (log10)
Sample 1 CV² (log10)
B
Sample 2 Fano (log10)
Sample 1 Fano (log10)
C
Rep 2 CV² (log10)
Rep 1 CV² (log10)
D
Rep 2 Fano (log10)
Rep 1 Fano (log10)
E
Median UMI Counts per Cell
count
40000
30000
20000
10000
0
DMSO_Jurkat_1
DMSO_Jurkat_2
IdU_Jurkat_1
IdU_Jurkat_2
DMSO_mESC_1
IdU_mESC_1

**Figure S3**

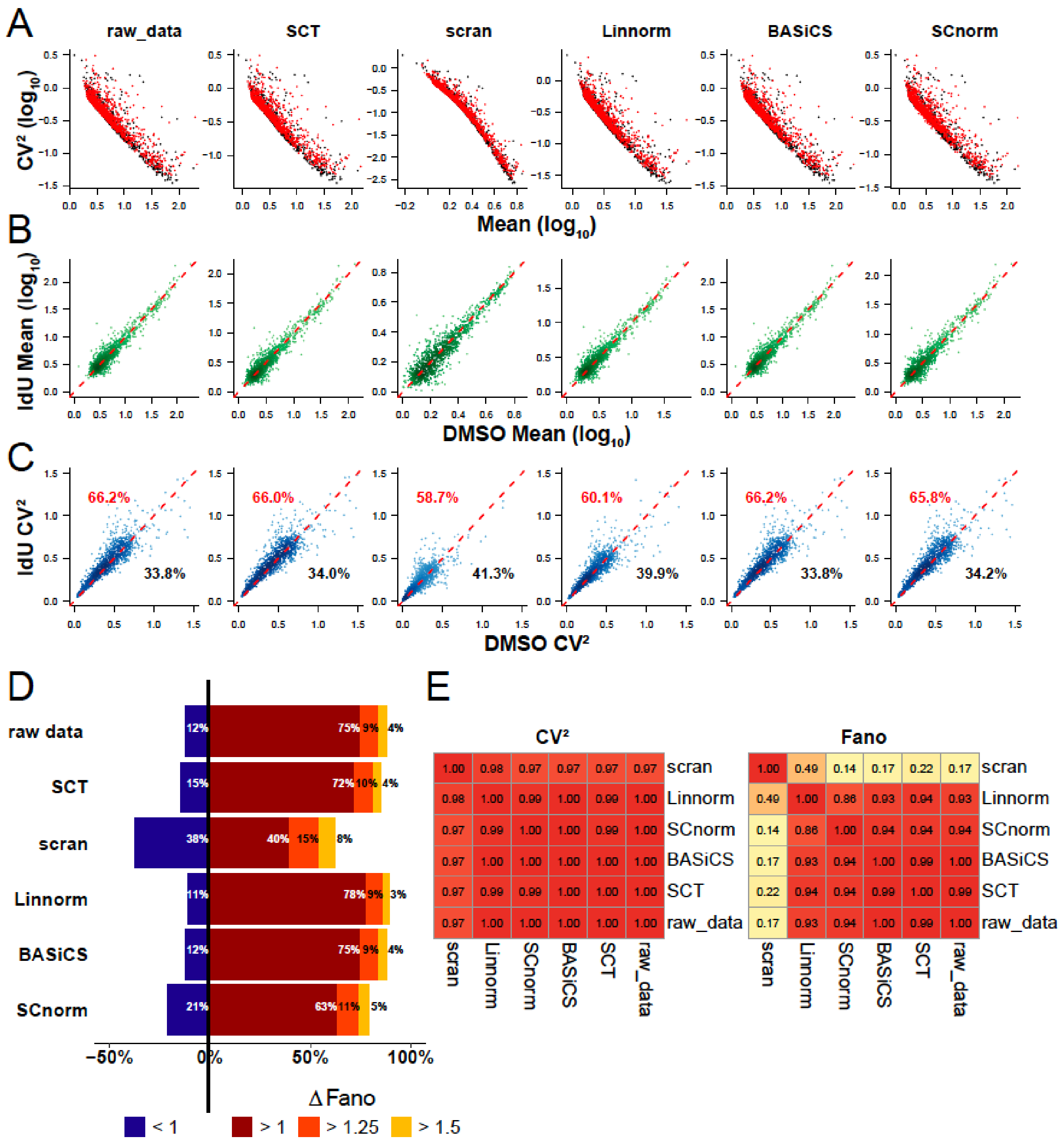
A
raw_data
SCT
scran
Linnorm
BASiCS
SCnorm
CV² (log10)
Mean (log10)
B
IdU Mean (log10)
DMSO Mean (log10)
C
IdU CV²
DMSO CV²
66.2%
33.8%
66.0%
34.0%
58.7%
41.3%
60.1%
39.9%
66.2%
33.8%
65.8%
34.2%
D
raw data
SCT
scran
Linnorm
BASiCS
SCnorm
12% 75% 9% 4%
15% 72% 10% 4%
38% 40% 15% 8%
11% 78% 9% 3%
12% 75% 9% 4%
21% 63% 11% 5%
-50%
0%
50%
100%
Δ Fano
< 1
> 1
> 1.25
> 1.5
E
CV²
1.00 0.98 0.97 0.97 0.97 0.97 scran
0.98 1.00 0.99 1.00 0.99 1.00 Linnorm
0.97 0.99 1.00 1.00 0.99 1.00 SCnorm
0.97 1.00 1.00 1.00 1.00 1.00 BASiCS
0.97 0.99 0.99 1.00 1.00 1.00 SCT
0.97 1.00 1.00 1.00 1.00 1.00 raw_data
scran Linnorm SCnorm BASiCS SCT raw_data
Fano
1.00 0.49 0.14 0.17 0.22 0.17 scran
0.49 1.00 0.86 0.93 0.94 0.93 Linnorm
0.14 0.86 1.00 0.94 0.94 0.94 SCnorm
0.17 0.93 0.94 1.00 0.99 1.00 BASiCS
0.22 0.94 0.94 0.99 1.00 0.99 SCT
0.17 0.93 0.94 1.00 0.99 1.00 raw_data
scran Linnorm SCnorm BASiCS SCT raw_data

**Figure S4**

**A**

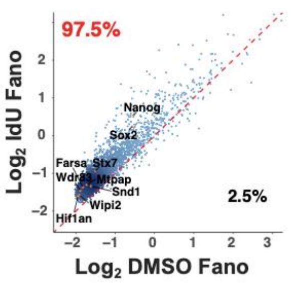


| Genes | Genomic location |
|---|---|
| *Nanog* | chr 6:122,707,568 |
| *Sox2* | chr 3:34, 650,405 |
| *Stx7* | chr 10:24,149,327 |
| *Farsa* | chr 8:84,856, 989 |
| *Wdr83* | chr 8:85,075, 035 |
| *Mtpap* | chr 18:4,375, 592 |
| *Hif1an* | chr 19:44,562,850 |
| *Wipi2* | chr 5:142,629 |
| *Snd1* | chr 6:28, 480,359 |

**B**

| Genes | *BASiCS* | | | *SCT* | | | *Scnorm* | | | *Scran* | | | *Linnorm* | | |
|---|---|---|---|---|---|---|---|---|---|---|---|---|---|---|---|
| | ΔMean | $\Delta CV^2$ | ΔFano | ΔMean | $\Delta CV^2$ | ΔFano | ΔMean | $\Delta CV^2$ | ΔFano | ΔMean | $\Delta CV^2$ | ΔFano | ΔMean | $\Delta CV^2$ | ΔFano |
| *Nanog* | 0.77 | 2.73 | 2.11 | 0.77 | 2.44 | 1.88 | 0.87 | 2.66 | 2.32 | 0.85 | 3.32 | 2.81 | 0.77 | 2.58 | 1.99 |
| *Sox2* | 0.59 | 2.96 | 1.75 | 0.59 | 2.56 | 1.50 | 0.55 | 2.66 | 1.46 | 0.76 | 5.13 | 3.89 | 0.63 | 2.76 | 1.72 |
| *Stx7* | 1.13 | 1.31 | 1.48 | 1.10 | 1.26 | 1.38 | 1.09 | 1.39 | 1.51 | 1.01 | 1.36 | 1.37 | 1.09 | 1.28 | 1.40 |
| *Farsa* | 1.00 | 1.37 | 1.37 | 1.01 | 1.22 | 1.22 | 1.03 | 1.39 | 1.42 | 0.98 | 1.47 | 1.45 | 0.99 | 1.34 | 1.32 |
| *Wdr83* | 0.93 | 1.46 | 1.35 | 0.89 | 1.27 | 1.14 | 0.83 | 1.42 | 1.17 | 0.91 | 1.55 | 1.41 | 0.94 | 1.43 | 1.34 |
| *Mtpap* | 1.01 | 1.33 | 1.34 | 0.99 | 1.30 | 1.28 | 1.07 | 1.44 | 1.54 | 0.95 | 1.49 | 1.42 | 1.00 | 1.38 | 1.38 |
| *Hif1an* | 0.94 | 1.34 | 1.26 | 0.91 | 1.30 | 1.18 | 0.77 | 1.19 | 0.92 | 0.89 | 1.33 | 1.19 | 0.95 | 1.31 | 1.25 |
| *Wipi2* | 1.02 | 1.24 | 1.26 | 1.00 | 1.13 | 1.13 | 0.86 | 1.24 | 1.07 | 0.98 | 1.37 | 1.34 | 1.01 | 1.23 | 1.25 |
| *Snd1* | 0.94 | 1.28 | 1.21 | 0.94 | 1.08 | 1.01 | 0.98 | 1.30 | 1.27 | 0.94 | 1.34 | 1.26 | 0.96 | 1.21 | 1.16 |

**C**

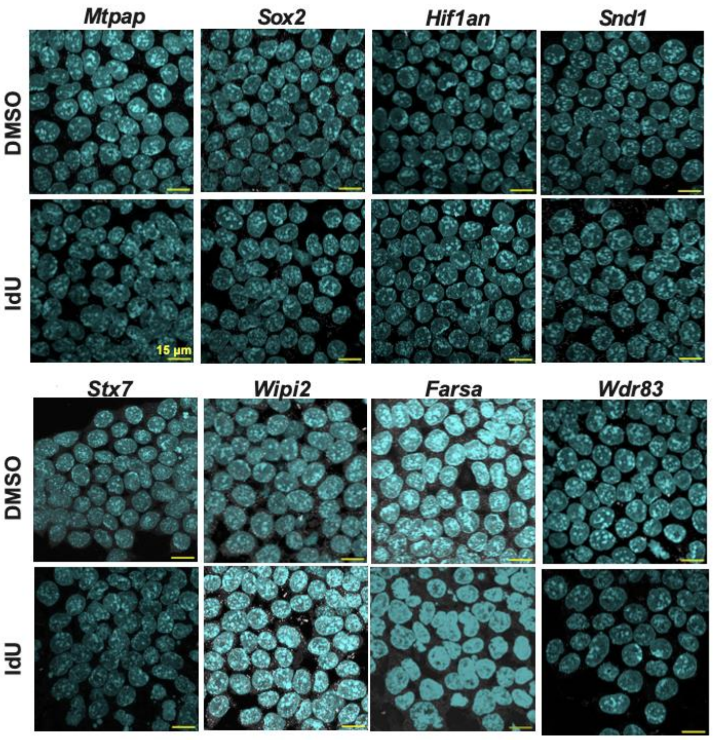


**D**

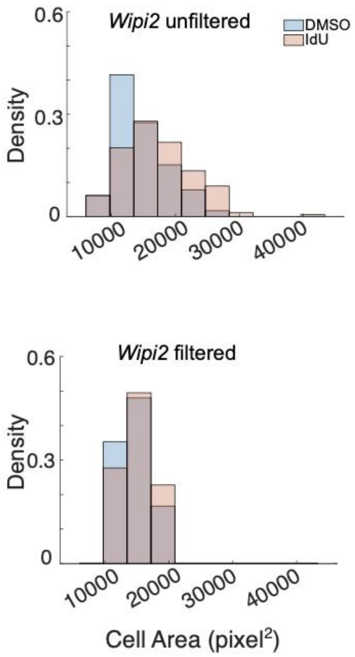

**Figure S5**

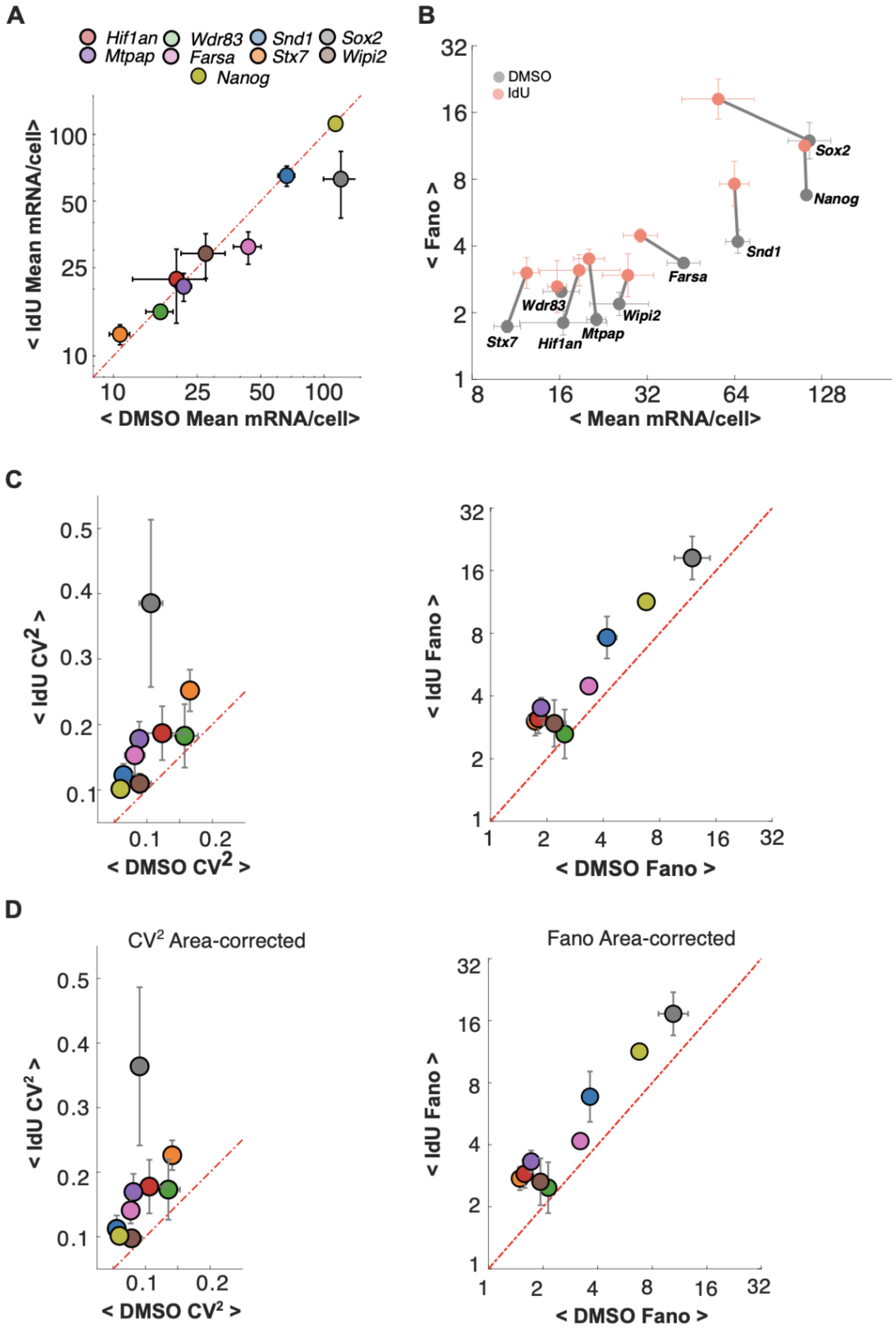
A
Hif1an
Wdr83
Snd1
Sox2
Mtpap
Farsa
Stx7
Wipi2
Nanog
< IdU Mean mRNA/cell>
< DMSO Mean mRNA/cell>
B
DMSO
IdU
< Fano >
< Mean mRNA/cell>
C
< IdU CV2 >
< DMSO CV2 >
< IdU Fano >
< DMSO Fano >
D
CV2 Area-corrected
Fano Area-corrected
< IdU CV2 >
< DMSO CV2 >
< IdU Fano >
< DMSO Fano >

**Figure S6**

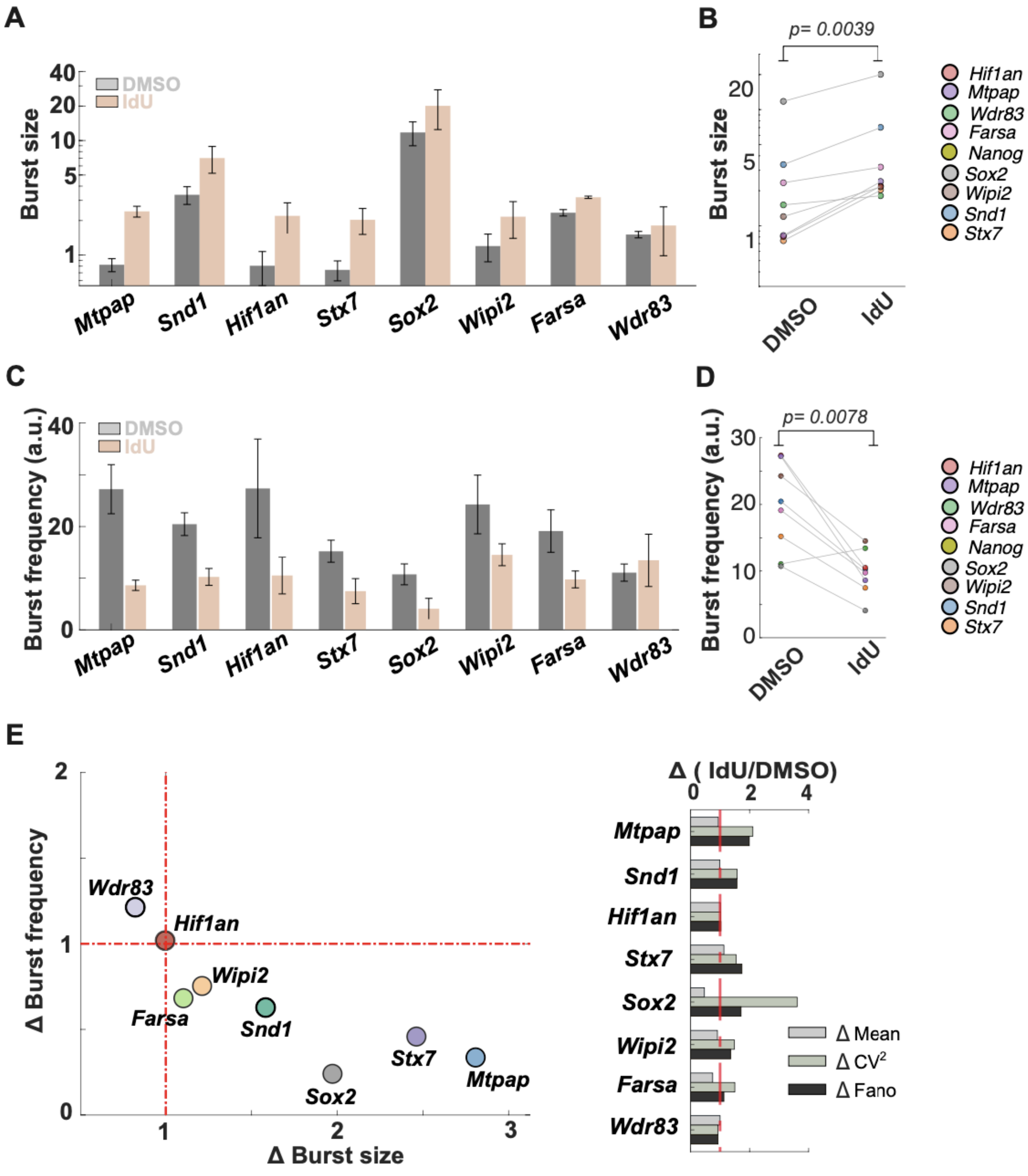
A
B
C
D
E
Burst size
DMSO
IdU
Mtpap
Snd1
Hif1an
Stx7
Sox2
Wipi2
Farsa
Wdr83
p= 0.0039
Nanog
Burst frequency (a.u.)
p= 0.0078
Δ Burst frequency
Δ Burst size
Δ ( IdU/DMSO)
Δ Mean
Δ CV2
Δ Fano
